# Polygenic selection underlies evolution of human brain structure and behavioral traits

**DOI:** 10.1101/164707

**Authors:** Evan R. Beiter, Ekaterina A. Khramtsova, Celia Van Der Merwe, Emile R. Chimusa, Corinne Simonti, Jason Stein, Paul Thompson, Simon E. Fisher, Dan J. Stein, John A. Capra, James A. Knowles, Barbara E. Stranger, Lea K. Davis

## Abstract

Seemingly paradoxical characteristics of psychiatric disorders, including moderate to high prevalence, reduced fecundity, and high heritability have motivated explanations for the persistence of common risk alleles for severe psychiatric phenotypes throughout human evolution. Proposed mechanisms include balancing selection, drift, and weak polygenic adaptation acting either directly, or indirectly through selection on correlated traits. While many mechanisms have been proposed, few have been empirically tested. Leveraging publicly available data of unprecedented sample size, we studied twenty-five traits (i.e., ten neuropsychiatric disorders, three personality traits, total intracranial volume, seven subcortical brain structure volume traits, and four complex traits without neuropsychiatric associations) for evidence of several different signatures of selection over a range of evolutionary time scales. Consistent with the largely polygenic architecture of neuropsychiatric traits, we found no enrichment of trait-associated single-nucleotide polymorphisms (SNPs) in regions of the genome that underwent classical selective sweeps (i.e., events which would have driven selected alleles to near fixation). However, we discovered that SNPs associated with some, but not all, behaviors and brain structure volumes are enriched in genomic regions under selection since divergence from Neanderthals ~600,000 years ago, and show further evidence for signatures of ancient and recent polygenic adaptation. Individual subcortical brain structure volumes demonstrate genome-wide evidence in support of a mosaic theory of brain evolution while total intracranial volume and height appear to share evolutionary constraints consistent with concerted evolution. We further characterized the biological processes potentially targeted by selection, through expression Quantitative Trait Locus (eQTL) and Gene Ontology (GO) enrichment analyses and found evidence for the role of regulatory functions among selected SNPs in immune and brain tissues. Taken together, our results suggest that alleles associated with neuropsychiatric, behavioral, and brain volume phenotypes have experienced both ancient and recent polygenic adaptation in human evolution, acting through neurodevelopmental and immune-mediated pathways.

## Introduction

Common (minor allele frequency [MAF] > 5%) single-nucleotide polymorphisms (SNPs) account for a substantial proportion of the inter-individual variation in psychiatric disorders and other brain-related phenotypes within human populations ^1–14^ Psychiatric disorders are often diagnosed before child-bearing years, and exhibit moderate to high prevalence worldwide, reduced fecundity ^15^, and high heritability ^16^. This raises a fundamental question of why risk alleles have persisted through the evolutionary history of our species ^17–19^. One proposed explanatory mechanism is balancing selection, which can increase frequencies of risk alleles through an advantage conferred to heterozygotes, as observed in sickle cell anemia and malarial resistance ^20^. Consistent with this hypothesis, siblings of individuals diagnosed with some neuropsychiatric disorders have significantly increased fecundity relative to the general population ^15^. An alternative explanation is that positive selection on genes influencing largely beneficial traits may have indirectly influenced psychiatric disorders through pleiotropic mechanisms ^21–23^. Another possibility that has been discussed in the literature is ancestral neutrality i.e., that reduced fecundity is only a modern phenomenon ^24^ For example, the “shaman theory” proposes that individuals with schizophrenia or other severe mental illness may have been considered endowed with mystical powers and thus not subject to modern stigma. ^24^ A more likely scenario, perhaps, is that women with developmental delay or psychiatric illness may have been (and continue to be) more vulnerable to rape ^25–27^ than women without psychiatric illness. Still others suggest that recent selection on genes involved in cognitive development may have over-selected the genes pre-disposing humans to mental disorders ^28, 29^. Finally, according to a theory known as polygenic mutation-selection balance, proposed in 2006 by Keller and Miller, psychiatric illness reflects a small mutational load dispersed across thousands of genes ^24^. Clearly, while many potential mechanisms have been hypothesized for risk variants maintaining a high allele frequency across evolutionary time, few have been empirically tested. Few genome-wide studies of adaptive selection have been reported for neuropsychiatric traits to date, thus motivating this study ^24, 28, 30^.

Paleobiological evidence indicates that the size of the human skull has expanded massively over the last 200,000 years, likely mirroring increases in brain size. It is thought that these structural changes in human brain size precipitated our uniquely human cognitive capabilities. The observed evolutionary increase in human skull size, however, is not sufficient to explain how individual brain structures have changed over evolutionary time. Our recent insights into the genetic contribution to total intracranial volume ^13, 31, 32^ and to subcortical brain structure volumes now provides an opportunity to look backwards in time and infer the history of brain structure during the evolution of humankind. Two prominent evolutionary theories, the “concerted” and “mosaic” theories of human brain evolution, may offer insight into the evolution of behavioral traits in our species. The concerted theory states that human brains evolved in concert with changes in body size and share developmental pathways that allometrically constrain the evolution of the brain ^33^. The mosaic theory, asserts that different brain structures are able to vary in size and evolve independently of one another^34–36^. Importantly, these theories are not mutually exclusive and there is evidence for both concerted and mosaic evolution of the brain across species ^37, 38^. Both theories carry implications for the evolutionary relationship between behavior and human brain development. For example, if evolution of the human brain is primarily a function of allometric scaling driven by genetic correlations with body size, behavior may have a more limited role in brain evolution. However, if behavioral traits are a target of selection, genetic correlation between those traits and individual brain structures could provide a mechanism to allow individual brain structures to adapt independently to a variety of evolutionary pressures. Finally, analysis of the signatures of selection across genetic variants associated with brain structure volumes may provide genetic and neurobiological insights into uniquely human capabilities.

Given recent advances in characterizing the genetic architecture of brain and behavioral phenotypes ^30, 32,39–52^ and analytic advances in detecting polygenic adaptation ^53, 54^, here we sought to systematically evaluate evidence for selection across a total of 25 complex traits including ten neuropsychiatric disorders, three personality traits, total intracranial volume, seven subcortical brain structure volume traits, and for comparison, four complex traits with no known neuropsychiatric associations (Supplementary Table 1). We tested summary statistics from publicly-available genome-wide association studies (GWAS) for multiple signatures of selection across different timescales of human evolutionary history, beginning with our divergence from the lineage that led to Neanderthals, estimated at ~600 kya. First, we examined trait-associated SNPs for enrichment in regions of the human genome that are depleted of Neanderthal alleles and hence thought to indicate positively selected regions since divergence from Neanderthals (i.e., Neanderthal selective sweep score, NSS) ^55^. Second, we tested for evidence of “soft selective sweeps” occurring up to 150 kya in which an adaptive mutation increases in frequency, but does not reach fixation, as indicated by global population differentiation (i.e., high global Fst,a metric of population differentiation due to genetic structure) ^56^. Third, we tested for classical “hard sweeps” occurring ~25-30 kya, in which a new adaptive mutation on a single haplotype quickly rises to fixation (or near fixation), leaving behind an identifiable signature in the form of reduced genetic diversity at the haplotype locus and longer than expected haplotype lengths (i.e., integrated haplotype scores, iHS) ^57–59^. Scans for signatures of hard sweeps have detected genes associated with traits such as lactase persistence ^60^, malaria resistance ^61^ and Glucose-6-phosphate dehydrogenase enzyme activity ^62^. Fourth, we employed an approach developed by Berg and Coop (2014) to detect weak positive selection that acts on standing variation (i.e., polygenic adaptation involving alleles at many different loci) and leaves behind a signature of increased covariance in trait-associated allele frequencies across 52 global populations compared to a model of genetic drift ^54, 63^. This approach is powered to detect polygenic adaptation approximately 30 kya. Fifth, we applied the Singleton Density Score (SDS) test, to investigate the role of very recent polygenic adaptation, occurring up to 2 kya. The SDS test allows detection of variants experiencing very recent positive selection because they have rapidly increased in frequency – giving the surrounding haplotypes little time to accumulate new mutations (i.e., singletons). Thus, for phenotypes evolving under recent positive polygenic adaptation, we expect to observe a reduced number of singleton events on the selected haplotype in reference to sequencing panels ^54^.

**Table 1.** Enrichment of trait-associated variants in genomic regions of extreme iHS, Fst, and S Score. The #SNPs denotes the number of trait-associated SNPs used in the analysis (Methods and Supplementary Figure 1). Enrichment p-values significant after multiple-testing correction (p<0.002) are denoted with an asterisk. 1: GWAS significance P < 5.0 × 10^−6^, 2: GWAS significance P < 5.0 × 10^−8^, 3: GWAS significance P < 5.0 × 10^−4^, 4: GWAS significance P < 5.0 × 10^−4^.

| Phenotypes | NSS-Score Enrichment |  | Extreme Population Differentiation |  |  | Extreme Integrated Haplotype Score Enrichment |  |  |
| --- | --- | --- | --- | --- | --- | --- | --- | --- |
| Neuropsychiatric and Behavioral Traits | #SNPs | Enrichment p-value ($S < -4.32$) | #SNPs | Enrichment p-value ($F_{st} > 0.30$) | Enrichment p-value ($F_{st} > 0.56$) | #SNPs | Enrichment p-value ($iHS > 2.0$) | Enrichment p-value ($iHS > 2.6$) |
| Attention Deficit and Hyperactivity Disorder | 2,319 | 0.544 | 1,036 | 0.048 | 0.624 | 514 | 0.552 | 0.760 |
| Alzheimer's Disease | 7,328 | 0.948 | 3,863 | 0.518 | 0.638 | 1,613 | 0.914 | 0.968 |
| Anorexia Nervosa | 8,200 | 0.934 | 4,247 | 0.868 | 0.168 | 1,512 | 0.944 | 0.714 |
| Generalized Anxiety Disorder | 5,564 | 0.896 | 2,504 | 0.934 | 0.678 | 1,317 | 0.974 | 0.368 |
| Autism Spectrum Disorder | 6,713 | 0.096 | 3,467 | 0.792 | 0.390 | 1,383 | 0.464 | 0.120 |
| Bipolar Disorder | 3,832 | 0.050 | 1,847 | <0.002* | 0.036 | 924 | 0.868 | 0.910 |
| Extraversion | 6,854 | 0.162 | 3,316 | 0.814 | 0.972 | 1,586 | 0.286 | 0.242 |
| Major Depressive Disorder | 2,412 | 0.044 | 1,162 | 0.146 | 0.632 | 595 | 0.966 | 0.890 |
| Neuroticism | 6,833 | <0.002* | 3,306 | 0.970 | 0.838 | 1,566 | 0.862 | 0.810 |
| Obsessive Compulsive Disorder | 7,475 | 0.230 | 3,930 | 1.000 | 0.994 | 1,624 | 0.904 | 0.960 |
| Schizophrenia | 16,200 | 0.002* | 8,759 | 0.378 | 0.860 | 3,845 | 0.974 | 0.972 |
| Subjective Well-Being | 3,858 | 0.436 | 1,806 | <0.002* | 0.234 | 1,011 | 0.898 | 0.928 |
| Tourette Syndrome | 7,917 | 0.532 | 4,246 | 0.878 | 0.750 | 1,694 | 0.594 | 0.774 |
| **Brain Structure Volumes** |  |  |  |  |  |  |  |  |
| Accumbens | 5,164 | 0.488 | 2,709 | 0.996 | 0.486 | 1,060 | 0.510 | 0.590 |
| Amygdala | 5,237 | 0.794 | 2,710 | 0.250 | 0.122 | 1,057 | 0.770 | 0.596 |
| Caudate Nucleus | 5,198 | 0.214 | 2,661 | 0.762 | 0.170 | 1,098 | 0.592 | 0.104 |
| Hippocampus | 6,621 | 0.594 | 3,462 | 0.944 | 0.966 | 1,366 | 0.998 | 0.906 |
| Total Intracranial Volume | 7,446 | 0.004 | 3,888 | 0.680 | 0.758 | 1,561 | 0.776 | 0.890 |
| Pallidum | 5,271 | 0.026 | 2,707 | 0.856 | 0.678 | 1,075 | 0.768 | 0.948 |
| Putamen | 5,064 | 0.766 | 2,662 | 0.640 | 0.504 | 1,097 | 0.942 | 0.778 |
| Thalamus | 5,204 | 0.288 | 2,649 | 0.902 | 0.220 | 1,064 | 0.508 | 0.776 |
| **Complex Traits** |  |  |  |  |  |  |  |  |
| BMI<sup>1</sup> | 306 | 0.032 | 179 | 0.260 | 0.124 | 107 | 0.968 | 0.578 |
| Height<sup>2</sup> | 1,449 | 0.108 | 838 | 0.150 | 0.394 | 405 | 0.480 | 0.972 |
| Inflammatory Bowel Disease<sup>3</sup> | 2,170 | 0.400 | 1,250 | 0.686 | 0.612 | 482 | 0.150 | 0.452 |
| Type 2 Diabetes<sup>4</sup> | 669 | 0.004 | 344 | 0.002* | 0.134 | 168 | 0.346 | 0.576 |

Typical effect sizes of common risk alleles for complex traits are small, so we hypothesized that there would be limited evidence of classical hard sweeps (typically targeting SNPs with large selection coefficients) on variants associated with these traits. Rather, we reasoned that if positive selection has played any role in maintaining risk allele frequencies for neuropsychiatric and brain structure volume traits, it would produce a distributed polygenic signal throughout the genome across many weakly selected variants. Furthermore, signatures of selection that differ across brain structure volumes would provide evidence in favor of mosaic brain evolution, while strong genetic correlations and consistent patterns of polygenic adaptation would provide evidence for concerted evolution.

Finally, recent studies have shown that several neuropsychiatric traits are genetically correlated with metabolic and immune-mediated phenotypes providing potential mechanisms through which pleiotropic evolutionary adaptation may influence trait-associated allele frequencies ^64, 65^. For example, immune genes show evidence of pleiotropy in early neuronal development among humans and animal models ^66^ and the immune system has been a target of positive selection as humans have colonized land masses across the world, experiencing various environmentally-mediated immune challenges ^63, 67^ Therefore, to characterize the potential biological targets of selection we analyzed the biological functions of trait-associated SNPs for those phenotypes showing evidence of polygenic adaptation through studies of expression Quantitative Trait Loci (eQTL) detected in relevant tissues, including brain regions (e.g., frontal lobe, cerebellum, etc.) and immune tissues (e.g., monocytes, CD4+ cells, and whole blood) and gene set enrichment. In summary, we systematically assessed multiple signatures of selection across a comprehensive set of brain and behavior phenotypes and further investigated potential biological mechanisms underlying significant evidence of adaptation.

## Methods

### Summary Statistics from Genome-Wide Association Studies

GWAS summary statistics for attention deficit hyperactivity disorder (ADHD)^1^, Alzheimer’s disease (ALZ)^2^, anorexia nervosa (AN)^3^, general anxiety disorder (GAD)^4^, autism spectrum disorder (ASD)^68^, bipolar disorder (BIP)^5^, major depressive disorder (MDD)^7^, obsessive compulsive disorder (OCD)^69^, Tourette Syndrome (TS)^8, 70^, schizophrenia (SCZ)^52^, extraversion (EXT)^6, 71^, neuroticism (NEO)^71, 72^, subjective wellbeing (SWB)^14^, type 2 diabetes (T2D)^10^, inflammatory bowel disease (IBD)^9^, height (HEI)^11^, body mass index (BMI)^12^, total intracranial volume (ICV)^32^, total intracranial volume controlled for height (ICV|height)^32^ and seven separate subcortical brain regions (the nucleus accumbens^13^, amygdala^13^, caudate nucleus^13^, pallidum^13^, putamen^13^, thalamus^13^ and hippocampus^31^) were obtained from consortium websites or directly from study authors (Supplementary Table 1). To protect research participants from re-identification in summary GWAS data, many consortia replace the true allele frequencies with allele frequencies drawn from the 1000 Genomes Project or HapMap, depending on author preference. We therefore used a common reference and determined the allele frequencies across phenotypes by calculating each within the 1000 Genomes European population. For each phenotype, the analysis was either restricted to European populations or included a very small number of non-European cases and controls that were meta-analyzed with a majority of European samples. Integrated haplotype scores (iHS) ^59^ and global Fst scores ^56^ were downloaded from the 1000 Genomes Selection Browser 1.0 ^73^. SNPs modestly associated with each trait (p < 5.0 × 10^−3^) were chosen for subsequent analysis. To test the robustness of the results against varying significance thresholds, we also tested a subset of SNPs exceeding a more stringent p-value threshold (p < 5.0 × 10^−4^). Trait-associated SNPs were clumped using an R^2^ threshold of 0.25, within a 500Kb window. Again, to determine the robustness of results against varying LD-pruning thresholds, we also generated a more stringently clumped set of SNPs for each phenotype (r^2^ < 0.10, 1000Kb window). We retained the most significantly associated SNP within each LD block determined by these parameters. These sets of clumped, nominally trait-associated SNPs were then utilized in enrichment and polygenic analyses.

### Enrichment of variants in regions of the genome under selection since Neanderthal divergence

Each set of trait-associated SNPs was annotated with Neanderthal selective sweep scores (NSS-scores) ^55^. Extreme negative NSS-scores (< −4.32, 5% tail of genome-wide distribution) indicate regions of the human genome that are depleted of derived Neanderthal alleles and hypothesized to have experienced selective sweeps in modern human ancestors occurring since the time of human divergence from Neanderthals ^55^. To test for enrichment of low NSS-scores among these top SNP associations, 500 sets of null matched SNPs were generated using SNPsnap^74^. For each phenotype, the 500 SNP sets were sampled without replacement from the European catalogue of 1000 Genomes SNPs. Randomly chosen SNPs were matched to the trait-associated SNPs by minor allele frequency (± 3%), gene density (± 50%), and distance to nearest gene (± 50 kb) for each phenotype. An empirical enrichment p-value was calculated representing the proportion of random, matched SNP sets in which the number of SNPs within the Neanderthal-derived depleted regions matched or exceeded the actual number of observed trait-associated SNPs within the Neanderthal-derived depleted regions for each phenotype.

### Enrichment of signatures of selective sweeps among trait associated SNPs

To test for enrichment of extreme iHS or Fst scores among these top SNP associations, 500 sets of randomly ascertained SNPs were generated using SNPsnap^74^. The same general enrichment approach used for NSS-scores was employed with one modification. SNPs of Neanderthal ancestry have demonstrated high iHS due to the extended LD caused by recent introgression ^75^. Therefore, to avoid confounding signatures of positive selection and introgression, SNPs located in regions of Neanderthal introgression were excluded from both trait-associated SNPs and the matching SNP sets ^76, 77^. Because integrated haplotype scores (i.e., our tests of “hard sweeps”) are based on detection of haplotype lengths that are longer than expected, we also did not match SNPs on LD parameters, to avoid overmatching the randomly chosen SNP sets.

Each trait-associated SNP set was then tested for enrichment of high iHS scores at two thresholds (|iHS| > 2.0 and |iHS| > 2.6) and extreme global population differentiation at two thresholds (F_st_ > 0.30 and F_st_ > 0.56), representing the most extreme 5% and 1%, respectively, of both scores across the human genome. An empirical enrichment p-value was calculated as the proportion of random matched SNP sets in which the percentage of extreme scores (i.e., iHS or F_st_, respectively) match or exceed the actual observed percentage of extreme scores in the set of trait-associated SNPs. (We summarize the enrichment analysis workflow in Supplementary Figure 1).

**Figure 1.**
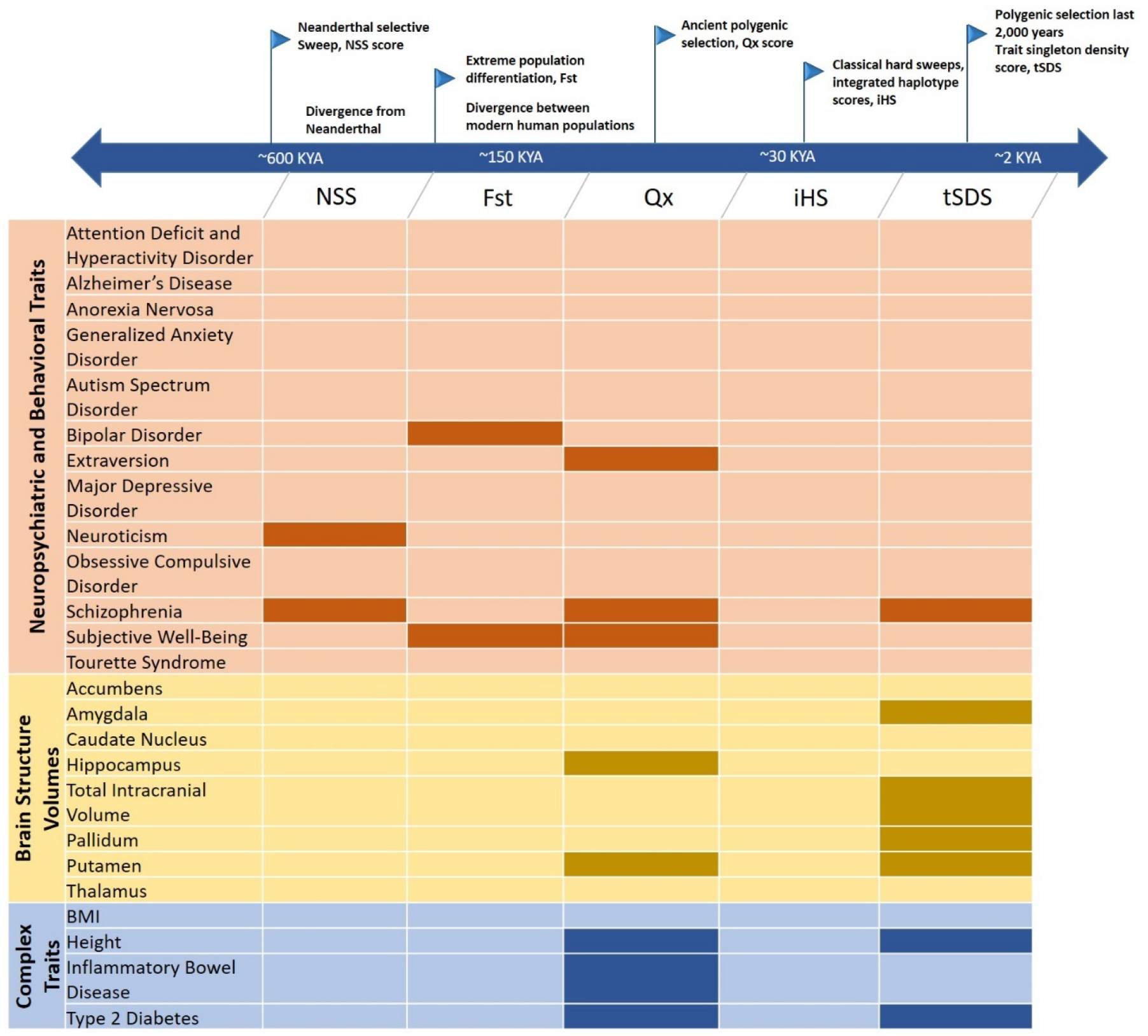
Summary of various ancient and recent signatures of polygenic adaptation analysis in neuropsychiatric and behavioral traits (orange), brain structure volumes (yellow), and complex traits (blue). The tested signatures are selection since Neanderthal divergence (Neanderthal selective sweep, NSS score), extreme population differentiation (fixation index, F_st_), ancient polygenic selection (Qx score), classical hard sweeps (integrated haplotype score, iHS), and polygenic selection in the past 2,000 years (trait singleton density score, tSDS). The boxes filled with a darker color for each of the tests represent significant signature of selection for a given phenotype after multiple testing correction for 25 phenotypes (p < 0.002) immune tissue or cell type: whole blood, monocytes, and CD4+ T cells, while the dark green represents enrichment in a combination of the three immune tissues. Light blue bars represent each brain tissue, while the dark blue represents enrichment in a combination of ten brain tissues, or all ten brain tissues minus cerebellum. The black dashed line represents a p-value of 0.05. The red dashed line represents the significant p-value threshold (0.004) after accounting for 13 tissues.

### Polygenic Adaptation Analysis

The polygenic adaptation detection methodology of Berg and Coop (2014) was used to test whether estimated genetic values (i.e., linear weighted sum of risk or trait-increasing alleles) exhibit more covariance among populations than expected due to genetic drift, tested empirically with a statistic termed the “Qx score”. For all twenty-one brain-related phenotypes, SNPs exceeding the nominal association threshold (p < 5.0 × 10^−3^) were included in the analysis (Table 2). For non-neuropsychiatric traits (i.e., height, body mass index, type 2 diabetes, and inflammatory bowel disease) in which publicly-available

**Table 2.** Enrichment of trait-associated variants in genomic regions experiencing polygenic adaptation. The selection test statistic (Qx) was computed as described in Berg and Coop (2014). Enrichment p-values significant after multiple-testing correction are denoted with an asterisk.

| Phenotypes | #SNPs | Qx | P(Qx) | #SNPs | Qx | P(Qx) |
| --- | --- | --- | --- | --- | --- | --- |
| **Neuropsychiatric Traits** | $P < 5.0 \times 10^{-3}$ | | | $P < 5.0 \times 10^{-4}$ | | |
| Alzheimer's Disease | 1,777 | 83.80 | 0.014 | 259 | 46.81 | 0.688 |
| Anorexia Nervosa | 1,196 | 68.96 | 0.112 | 166 | 51.41 | 0.556 |
| Generalized Anxiety Disorder | 1,420 | 66.41 | 0.141 | 183 | 59.21 | 0.235 |
| Autism Spectrum Disorder | 1,149 | 58.02 | 0.334 | 138 | 42.69 | 0.833 |
| Bipolar Disorder | 2,634 | 57.26 | 0.310 | 449 | 54.30 | 0.356 |
| Extraversion | 1,485 | 88.04 | 0.001* | 196 | 49.71 | 0.545 |
| Major Depressive Disorder | 1,806 | 63.31 | 0.146 | 224 | 70.11 | 0.065 |
| Neuroticism | 1,617 | 77.52 | 0.156 | 205 | 75.69 | 0.029 |
| Obsessive Compulsive Disorder | 1,198 | 70.33 | 0.082 | 172 | 39.18 | 0.905 |
| Schizophrenia | 3,307 | 208.36 | <0.001* | 1,029 | 101.30 | <0.001* |
| Subjective Well-Being | 2,717 | 113.90 | <0.001* | 434 | 74.87 | 0.029 |
| Tourette Syndrome | 1,441 | 73.51 | 0.082 | 209 | 42.79 | 0.828 |
| **Brain Structure Volumes** | $P < 5.0 \times 10^{-3}$ | | | $P < 5.0 \times 10^{-4}$ | | |
| Accumbens | 1,152 | 61.61 | 0.203 | 134 | 63.86 | 0.162 |
| Amygdala | 1,164 | 55.05 | 0.378 | 137 | 53.34 | 0.470 |
| Caudate Nucleus | 1,210 | 48.33 | 0.626 | 153 | 56.18 | 0.348 |
| Hippocampus | 1,237 | 108.98 | <0.001* | 177 | 79.65 | 0.010 |
| Intracranial Volume | 1,249 | 54.37 | 0.386 | 175 | 50.79 | 0.526 |
| Pallidum | 1,183 | 52.76 | 0.559 | 176 | 48.78 | 0.625 |
| Putamen | 1,203 | 115.76 | <0.001* | 155 | 72.69 | 0.029 |
| Thalamus | 1,123 | 70.98 | 0.088 | 153 | 57.51 | 0.288 |
| **Complex Traits** | $P < 5.0 \times 10^{-4}$ | | | $P < 5.0 \times 10^{-6}$ | | |
| Inflammatory Bowel Disease | 423 | 101.13 | <0.001* | 81 | 77.25 | 0.017 |
| Type 2 Diabetes | 478 | 117.77 | <0.001* | 42 | 50.61 | 0.525 |
| | $P < 5.0 \times 10^{-6}$ | | | $P < 5.0 \times 10^{-8}$ | | |
| BMI | 246 | 78.20 | 0.010 | 97 | 82.61 | 0.003 |
| Height | 2,002 | 303.29 | <0.001* | 1,087 | 209.27 | <0.001* |

GWAS results report analysis of hundreds of thousands of individuals, we tested more stringent association thresholds (p < 5.0 × 10^−4^, p < 5.0 × 10^−6^, or p < 5.0 × 10^−8^) due to the computational burden of including nominally associated SNPs from these well-powered studies. For each phenotype, in each of 52 world populations, the genetic value was calculated as the linear sum of the number of risk (or trait-increasing) alleles in the population weighted by the effect size from the discovery GWAS (Supplementary Table 1)

In brief, Qx scores quantify the variance and covariance of allele frequencies across 52 human populations to detect the over-dispersion of risk alleles across populations in comparison to an empirical model of neutral drift taking into account shared ancestry. To create the model of neutral drift, a null distribution was created for each set of trait-associated SNPs, composed of 1000 sets of SNPs matched to the trait-associated SNPs by MAF (2% bins) and B-value, a metric of neutral genetic drift (100 bins) ^78^. Any SNP that was fixed in all 52 populations (or in the French European population) or was not part of the Human Genome Diversity Panel dataset was excluded from analysis (Supplementary Figure 1). The significance of the trait-associated SNP Qx statistic was determined empirically as described above.

To ensure that the model of genetic drift (conditioned based on ancestral allele frequencies) is unbiased, we compared the genetic values for each of the 52 populations to their *F*_3_ statistic described by Patterson et al (2012)^79^ in the form of:

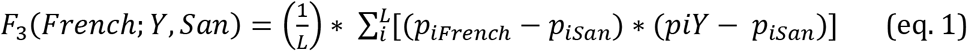

in which *p_i_* is the allele frequency at locus *i* and L = 20,000 randomly ascertained SNPs (Equation 1). No phenotypes demonstrated any correlation between genetic value and the *F_3_* statistic indicating that our results cannot be explained by a biased modeling of genetic drift (Supplementary Table 2; Supplementary Figure 3).

### Singleton Density Score Analysis

The singleton density score approach is based on the assumption that alleles under positive selection will rise in frequency faster than their haplotypes can accumulate new mutations. This results in an observable decrease in the density of singleton events on the selected haplotype measured by sequencing of large reference populations^54^. Scores for each SNP can be calculated in which haplotypes are annotated as ancestral and derived (i.e., singleton density scores or SDS) or calculated with respect to risk (or trait-increasing) and protective (or trait-decreasing) alleles (i.e., trait-singleton density scores or tSDS). To test whether polygenic positive selection is acting on risk or protective (trait-increasing or trait-decreasing) alleles we tested the global Spearman correlation between tSDS and strength of association from the GWAS. As in Field et al., 2016, each complete set of GWAS summary statistics was divided into 100 blocks of contiguous SNPs and a block jackknife resampling approach (i.e., leave-one-block-out in 100 iterations) was used to calculate the standard error while accounting for LD. A positive tSDS implies very recent selection (~2,000 years) on the risk (trait-increasing) allele while negative tSDS suggests recent selection on the protective (trait-decreasing) allele.

### Assessment of significance

For each of the five signatures selection, empirical assessments of the significance of the test statistic observation were conducted as described in respective methods sections above. In each case, we accounted for the number of phenotypes tested and considered the result significant if it met or exceeded the Bonferroni correction for all 25 phenotypes tested (p < 0.002).

### Enrichment of expression quantitative trait loci in brain and immune tissues

To investigate the biological functions of the trait-associated SNPs for the phenotypes with evidence of polygenic selection, we annotated SNPs with expression quantitative trait loci (eQTLs) information. We used previously published significant cis-eQTL results derived from 10 brain regions and whole blood ^80^ (https://www.gtexportal.org/home/), as well as CD4+ T cells (adaptive immunity) and CD14+ monocytes (innate immunity) ^81^.

To assess eQTL enrichment, specifically, to test for an enrichment for a gene regulatory role for trait-associated SNPs, we quantified the enrichment of the number of eQTL target genes (eGenes) associated with trait-associated SNPs compared to random, matched SNPs. One thousand randomly ascertained SNP sets were generated for each phenotype using SNPsnap ^74^. Matching sets were sampled without replacement from the European catalogue of 1000 Genomes SNPs, and matched for physical distance within 500 kb, MAF (± 3%), gene density (± 50%), distance to nearest gene (± 100 kb), and LD buddies (± 50%) at r^2^=0.8. Trait-associated SNPs and matching SNP sets were annotated as cis-eQTLs, and the genes they are associated with, as eGenes, in various tissues. The number of eGenes in each matched set yielded an empirical distribution. The enrichment P-value was calculated as the proportion of randomized sets in which the eGene count matches or exceeds the actual observed count in the list of the trait-associated SNPs. If several SNPs implicated the same eGene in a tissue, the eGene was counted once. The enrichment was considered significant if it met Bonferroni multiple testing correction threshold p<0.004 (i.e., 0.05/13 tissues investigated). Due to a high degree of eQTL sharing across tissues ^82^, when enrichment was assessed in combined brain tissues and combined immune tissues, only the set of unique eGenes was counted. To exclude the possibility of eQTL enrichment overestimation due to the gene-rich MHC region, we performed eQTL enrichment analysis both including and excluding SNPs in the HLA region.

To evaluate underlying biological pathways that may be regulated by the eQTLs identified in the brain and immune tissues, we applied Gene Ontology (GO) enrichment analysis as implemented in Gene Set Enrichment Analysis (GSEA) ^83, 84^ using the eGene lists as the input genes, and the whole genome as the background.

### Proportion of ancestral and derived alleles among risk conferring SNPs

The SNP ancestral allele annotations were derived from the Ensembl 59 comparative species alignment or were obtained from dbSNP. For each phenotype, we determined the proportion of risk alleles (p< 5.0 × 10^−3^) that were ancestral or derived. Previous work has shown that derived alleles are more often minor alleles (< 50% allele frequency) and more often associated with risk than ancestral alleles ^85^. For each phenotype, we investigated the proportion of derived risk alleles across minor allele frequency bins. To this end, the risk alleles were categorized into 5 MAF bins, (>0.05-0.1, >0.1-0.2, >0.2-0.3, >0.3-0.4, >0.4-0.5), and we independently computed the fractions of derived alleles in each bin.

## Results

### Variants associated with several brain phenotypes are enriched in regions of the genome under selection since Neanderthal divergence

SNPs nominally associated with schizophrenia (pscz = 0.002) and neuroticism (pneu < 0.002), were significantly enriched in regions hypothesized to have experienced positive selection early in modern human evolution (i.e., low NSS-score SNPs) compared to randomly ascertained matched SNPs (Methods and Table 1). SNPs nominally associated with total intracranial volume (p_ICV_ = 0.004) and Type 2 Diabetes (pt2d = 0.004) trended toward significant enrichment, however the remaining phenotypes demonstrated no enrichment, and none of the phenotypes demonstrated depletion.

### No evidence for strong selective sweeps among SNPs associated with brain and behavior phenotypes

None of the neuropsychiatric trait-associated SNPs exhibited enrichment of extreme iHS scores characteristic of hard sweeps of positive selection (Table 1). No phenotype demonstrated consistent, robust evidence for population differentiation. While SNPs associated with bipolar disorder and subjective well-being, respectively, showed significant evidence (pbpd < 0.002, pswb < 0.002) of enrichment for global population differentiation at the less extreme F_st_ threshold (F_st_ > 0.30, representing the top 5% of differentiated SNPs in the genome), they demonstrated no enrichment at the more stringent F_st_ threshold (F_st_ > 0.56, representing the top 1% of differentiated SNPs in the genome). All analyses, including enrichment of extreme F_st_ scores, can be influenced by residual population structure that may remain even after extensive QC procedures are performed. To test whether our results were influenced by residual population stratification, we calculated LD-score regression intercepts and lambda values (Supplementary Table 1), which reflect the contribution of confounding bias due to cryptic relatedness and residual population stratification (i.e., scores close to 1 indicate minimal residual confounding). No significant evidence of residual population stratification or cryptic relatedness was observed.

### Schizophrenia associated loci exhibit evidence of ancient polygenic adaptation

All traits were evaluated for evidence of ancient polygenic adaptation (approximately 30 kya) by testing each set of nominally associated SNPs for excess genetic covariance, after accounting for background selection, compared to a null model of genetic drift. Schizophrenia (Qx = 208.36, pscz < 0.001) was the only neuropsychiatric disorder that provided statistically significant evidence of ancient polygenic adaptation (Figure 1, Table 2, Supplementary Figure 2). Among the personality traits, extraversion (Qx = 88.04, pext < 0.001) and subjective well-being (Qx = 113.90, pswb < 0.001), but not neuroticism (Qx = 77.52, pneu = 0.156) demonstrated significant evidence of ancient polygenic adaptation (Figure 1). Moreover, among the brain structure volume traits, hippocampus (Qx = 108.98, phip < 0.001) and putamen (Qx = 115.76, pput < 0.001) demonstrated significant evidence of ancient selection (Figure 1, Table 2, Supplementary Figure 2). For comparison, we also examined common complex phenotypes with no known neuropsychiatric component. Inflammatory bowel disease (Qx = 101.13, pibd < 0.001), type 2 diabetes (Qx = 117.77, pt2d < 0.001), and height (Qx = 303.29, phei < 0.001) demonstrated significant evidence of polygenic adaptation (Table 2, Figure 1), consistent with previous reports ^53, 86–88^.

**Figure 2.**
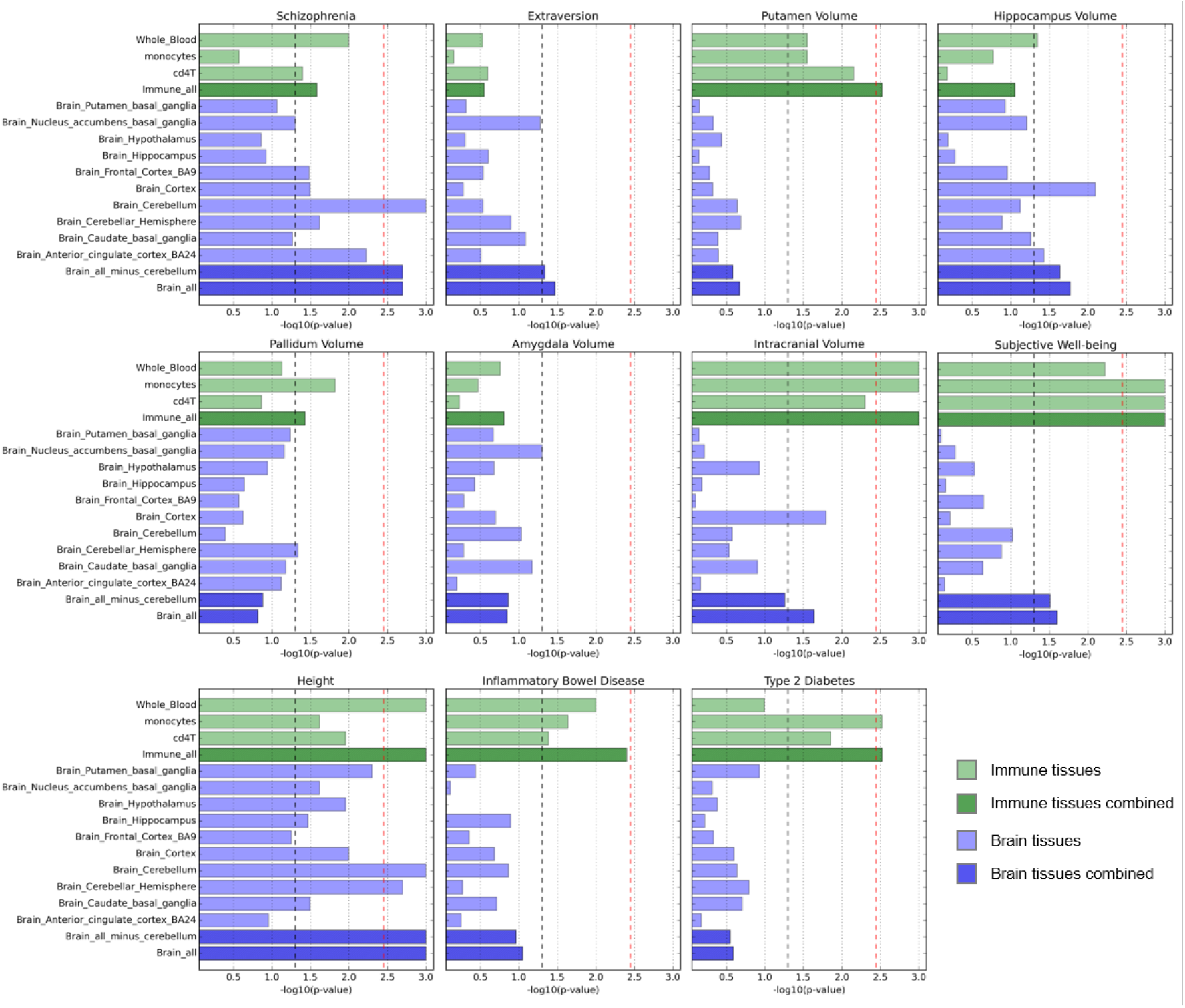
eQTL enrichment in the brain and immune tissues for eleven phenotypes (excluding SNPs in the HLA region) exhibiting evidence of polygenic selection (significant Qx or tSDS after multiple testing correction). Light green bars represent each immune tissue or cell type: whole blood, monocytes, and CD4+ T cells, while the dark green represents enrichment in a combination of the three immune tissues. Light blue bars represent each brain tissue, while the dark blue represents enrichment in a combination of ten brain tissues, or all ten brain tissues minus cerebellum. The black dashed line represents a p-value of 0.05. The red dashed line represents the significant p-value threshold (0.004) after accounting for 13 tissues.

When a more stringent GWAS association threshold was applied (GWAS p < 5.0 × 10^−4^), the number of SNPs included in each selection analysis dropped, resulting in attenuation of the polygenic adaptation signal for extraversion (Qx = 49.71, pext = 0.545), subjective well-being (Qx = 74.87, pswb = 0.029), hippocampus volume (Qx = 79.65, phip = 0.01), putamen volume (Qx = 72.69, pput = 0.029), inflammatory bowel disease (Qx = 77.25, pibd = 0.017), and type 2 diabetes (Qx = 50.61, pt2d = 0.525). Results for schizophrenia and height, both of which suffered less SNP attrition at the more stringent GWAS p-value, were robust (Qx = 101.30, pscz < 0.001; Qx = 209.27, phei < 0.001) to the more stringent association threshold, likely due to the power of the original GWAS (Table 2).

Because the null model of genetic drift is computed with respect to the derived and ancestral alleles (not necessarily the trait-associated alleles), the direction of selection with respect to risk can be difficult to determine. Most phenotypes tested exhibited a higher proportion of derived risk or trait-increasing alleles (for quantitative traits); however, there were six phenotypes with a slightly increased proportion of ancestral risk (or trait-increasing) alleles – namely major depressive disorder, attention deficit hyperactivity disorder, body mass index, type 2 diabetes, height and bipolar disorder (Supplementary Figure 7).

### Brain structure volumes exhibit evidence for very recent mosaic polygenic adaptation

Several brain structure volumes exhibited evidence of very recent polygenic adaptation (approximately 2kya). Total intracranial volume exhibited significant positively shifted tSDS (Z = 6.23, p_ICV_ = 4.79 × 10^−10^) and also when intracranial volume was controlled for height (Z = 3.35, picv|height = 7.95 × 10^−4^), while amygdala (Z = −3.50, pamy =4.69 × 10^−4^), pallidum (Z = −3.40, ppal = 6.80 × 10^−4^), and putamen (Z = −4.93, pput = 8.15 × 10^−7^) exhibited significant negatively shifted tSDS (Supplementary Figure 5) indicating positive selection for *increased* total intracranial volume, but *decreased* volume of subcortical brain structures.

### Schizophrenia-protective alleles exhibit evidence for very recent polygenic adaptation

Schizophrenia demonstrated significant negatively shifted tSDS (Z = −3.07; p =0.002) indicating very recent positive selection on schizophrenia-protective alleles (Figure 1, Table 3, Supplementary Figure 4). Height exhibited significant positively shifted tSDS (Z = 17.14, phei 6.90 × 10^−66^) indicating very recent positive selection on height-increasing alleles; while T2D exhibited significant negative shifted tSDS (Z = – 6.68, pt2D =2.34 × 10^−11^) indicating positive selection on T2D-protective alleles, consistent with prior reports ^54^.

**Table 3.**
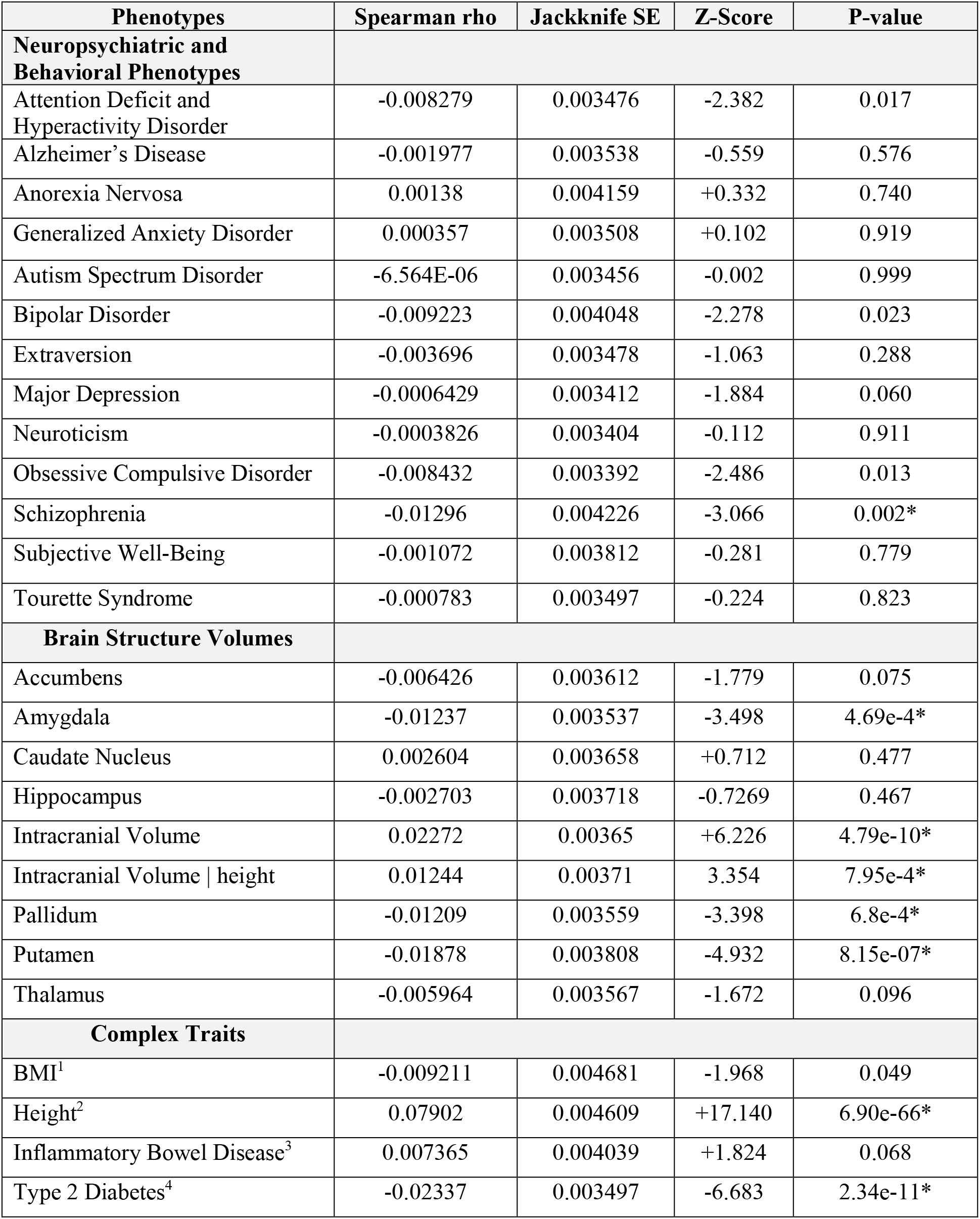
Recent positive selection on trait-associated variant using Singleton Density Score (SDS) analysis. The mean tSDS for the top 1,000 associated SNPs are shown: a negative value indicates positive selection on the trait-increasing allele, a positive value indicates positive selection on the trait-decreasing allele. The p-value for a two-tailed T-test represents deviation from the expected mean of 0. Results significant after multiple-testing correction are denoted with an asterisk. 1 GWAS significance P < 5.0 × 10^−6^, 2 GWAS significance P < 5.0 × 10^−8^, 3 GWAS significance P < 5.0 × 10^−4^, 4 GWAS significance P < 5.0 × 10^−4^.

### Brain and immune eQTLs are enriched among schizophrenia and brain structure volume-associated variants

For any phenotype exhibiting evidence of polygenic selection (significant Qx or tSDS after multiple testing correction), we tested whether trait-associated alleles are enriched for regulatory function in either brain or immune tissues. Given the fact that the polygenic selection scores are unable to robustly distinguish individual SNPs or alleles that have experienced selection, we examined in bulk the trait-associated SNPs that contributed to signatures of selection for these phenotypes. We expand on previous studies characterizing eQTL enrichment among neuropsychiatric trait-associated variants^8, 89^, by characterizing the functions of the implicated eGenes. We chose to focus on brain and immune eQTLs because both tissues have been previously implicated in neuropsychiatric and brain-related phenotypes and both systems have demonstrated adaptation throughout human evolution ^49, 90–92^. We performed cis-eQTL enrichment analysis using multiple brain tissues (putamen, nucleus accumbens, hypothalamus, hippocampus, frontal cortex, cortex, cerebellum, cerebellar hemisphere, caudate, and anterior cingulate cortex) and immune tissues (whole blood, CD4+ T cells, and CD14+ monocytes) ^80, 81^. Specifically, we tested for an enrichment for a gene regulatory role for trait-associated SNPs, as quantified by an enrichment of the number of eQTL target genes (eGenes) associated with trait-associated SNPs compared to random, matched SNPs. (Methods). This differs from previous eQTL enrichment analyses for neuropsychiatric phenotypes^8, 89^, in which the enrichment was based on whether a SNP was or was not an eQTL (ie., not accounting for the number of genes it may be associated with).

Among the brain and behavioral phenotypes with Bonferroni-corrected significant evidence for ancient or recent polygenic adaptation, only schizophrenia was significantly enriched for brain eQTLs (p_brain_all_ = 0.002, p_cerebellum_ < 0.001). eQTLs from immune cell types were strongly enriched among SNPs associated with subjective well-being (p_immune_all_ < 0.001, p_monocytes_ < 0.001, p_CD4+cells_ <0.001), intracranial volume (p_immune_all_ < 0.001, p_monocytes_ = 0.001, p_CD4+cells_ =0.005, p_whole blood_ <0.001), and putamen volume (p_immune_all_=0.003). Hippocampus, pallidum and amygdala volumes and extraversion showed no significant enrichment for brain or immune eQTLs (Figure 2, Supplementary Table 3). Including SNPs located in the HLA region did not significantly affect the results (Supplementary Figure 5, Supplementary Table 3). Additionally, we replicated previous findings demonstrating enrichment of brain eQTLs among BMI-associated SNPs (p_brain_all_ < 0.001), and enrichment of immune tissue eQTLs among SNPs associated with height (p_immune_all_<0.001), T2D (p_immune_all_=0.003), and IBD (p_immune_all_=0.004).

To characterize the sets of genes implicated by brain or immune eQTLs (i.e., eGenes, Supplementary Table 4), we performed GO term enrichment analysis on the eGenes derived from each phenotype that showed a significant tissue-specific enrichment (Supplementary Table 5). Here we report the top two significant molecular processes and biological processes for each phenotype-tissue pair. For schizophrenia, the brain eGenes showed the strongest enrichment for phosphate binding (q=1.24 × 10^−5^) and protein phosphatase binding (q=1.36 × 10^−04^) molecular functions, as well as negative regulation of molecular function (q=1.36 × 10^−04^) and establishment of localization in cell (q=2.19 × 10^−4^) biological processes. Intracranial volume immune eGenes showed the strongest enrichment for phosphoric ester hydrolase activity (q=5.54 × 10^−6^) and hydrolase activity acting on ester bonds (q=5.54 × 10^−6^) molecular functions, as well as dephosphorylation (q=4.68 × 10^−6^) and phosphate containing compound metabolic processes (q=3.20 × 10^−5^) biological processes. Subjective well-being immune eGenes showed the strongest enrichment for adenyl nucleotide binding (q=4.78 × 10^−6^) and ribonucleotide binding (q=1.41 × 10^−5^) molecular functions, as well as small molecule metabolic process (q=1.62 × 10^−9^) and organonitrogen compound metabolic process (q=1.22 × 10^−7^).

### Proportion of ancestral and derived alleles among risk conferring SNPs

For common risk alleles (MAF > 5%), the fraction that were also derived alleles was close to 50% for each phenotype tested (Supplementary Figure 6). For some phenotypes, such as eating disorders, Tourette Syndrome, Alzheimer’s disease, obsessive-compulsive disorder, and Type 2 diabetes, > 60% of risk alleles with MAF <5% were also derived alleles (Supplementary Figure 6).

## Discussion

The largely polygenic architecture of neuropsychiatric and other brain related phenotypes has been well established by studies demonstrating that the individual effect size of each common SNP is small, but cumulatively these SNPs account for a significant proportion of the estimated trait heritability ^51, 93–96^. For most phenotypes, a small proportion of cases appear to result from *de novo* mutations of large effect which disrupt the coding or regulatory sequences of critical brain genes ^90, 97–99^. The high penetrance and very low frequency of these variants in the general population suggest that they are likely targets of negative selection. In contrast, little is known about the history of selection on common genetic variation that confers risk for neuropsychiatric phenotypes. Here we present a systematic study of genetic evidence for strong and weak positive selection across a catalogue of twenty-five phenotypes including neuropsychiatric traits, personality traits, and brain structure volumes.

Consistent with the polygenic architecture of common variation underlying complex traits, we find little evidence of widespread global population differentiation and no evidence of strong selective sweeps among trait-associated SNPs (Table 1). While prior studies have reported significant genetic correlation between polygenic risk across ancestries ^100^, we cannot rule out the possibility that SNPs associated with these phenotypes in non-European ancestries may demonstrate greater population differentiation. Nevertheless, in contrast to autoimmune phenotypes ^87^, there is currently no evidence of strong positive selection acting on common risk alleles for neuropsychiatric disorders, personality traits, or brain structure volumes.

We observed a significant enrichment of trait-associated SNPs for schizophrenia and neuroticism within regions of the genome depleted of derived Neanderthal alleles, and nearly significant enrichment for total intracranial volume, consistent with the hypothesis that these genomic regions harbor variation fundamental to the development of the modern human brain ^30^. Importantly, this enrichment analysis focuses on regional genomic annotations, as opposed to allelic annotations thus we can make no claim regarding risk or protective alleles for any trait. Furthermore, we found no evidence of enrichment in these regions in some of the most well-powered traits we studied – including height and BMI – indicating that the findings in schizophrenia, neuroticism, and total intracranial volume were not solely a function of power achieved in the discovery GWAS.

Ancient polygenic adaptation (~30 kya), which is manifest in small but coordinated shifts in trait-associated frequencies, was observed among neuropsychiatric phenotypes (schizophrenia), personality traits (extraversion and subjective well-being), volumes of subcortical brain structures (putamen and hippocampus), autoimmune disorders (inflammatory bowel disease), height, and type 2 diabetes (Table 2). Loci associated with these traits demonstrated significant evidence of shifts in allele frequency consistent with a positive selection model. However, a limitation of this approach is that the null model of neutral drift is computed with respect to ancestral and derived allelic annotations, thus complicating the interpretation of whether ancient positive selection has acted on risk/trait-increasing alleles or protective/trait-decreasing alleles.

We employed a directional test of selection (tSDS) which overcomes this limitation, albeit covering a narrower evolutionary timescale. The tSDS approach is powered to detect selection that has been active during the past ~2,000 years – which makes it difficult to determine with certainty whether the observed direction of selection also reflects more ancient selection pressures ^54^ Using this test, we reproduced significant findings of positive polygenic selection acting on height-increasing alleles, T2D-protective alleles, and schizophrenia-protective alleles. Moreover, our tSDS analysis revealed strong evidence of recent polygenic selection acting on volumes of the amygdala, pallidum and putamen, as well as on total intracranial volume, and total intracranial volume adjusted for height.

The results provide evidence for both the mosaic and concerted theories of brain evolution. Specifically, we find that recent positive polygenic selection has strongly favored total intracranial-volume *increasing* alleles, which closely mirrors the adaptation signal observed for height*-increasing* alleles. A significant genetic correlation between height and total ICV has also been reported (Rg = 0.241, p = 2.4 × 10^−10^)^32^. In the analysis of ICV adjusted for height, the strength of the signal for polygenic adaptation was greatly attenuated (Table 3) but remained significant even after accounting for substantial evolutionarily constraint related to overall body size. Integration of eQTLs from the GTEx project further characterizes this relationship revealing that approximately 30% of the brain-tissue eGenes targeted by ICV-associated SNPs are also targeted by height-associated SNPs, suggesting a shared set of genes influencing both phenotypes and experiencing positive polygenic selection.

Among subcortical brain volume traits, we observed signatures of ancient and recent adaptation in some (e.g., hippocampus, putamen, pallidum, amygdala), but not all (e.g., nucleus accumbens, caudate nucleus, thalamus) brain structures (Figure 1). The differential pattern of signatures of selection suggest that individual brain structures have been able to evolve independently of each other, consistent with a mosaic theory of brain evolution ^34–36, 101^. Furthermore, our results indicate very recent positive selection acting on total intracranial *volume-increasing* alleles, but putamen, amygdala and pallidum volume-*decreasing* alleles (Supplementary Figure 5). While increases in some regions could be genetically correlated to decreases in other regions, GWAS findings from Hibar et al. (2015) also confirm that despite high heritability, the genetic correlations between the structures are low. Thus, in the context of previous findings, our results provide direct genetic evidence for a model of selection that has differentially influenced both volume-increasing and volume-decreasing alleles, perhaps providing opportunities for modern human-specific neocortex enlargement over time. If correct, falsifiable hypotheses also emerge from these findings; for example, cortical surface area-increasing alleles should demonstrate evidence of positive selection during both recent and ancient evolutionary history. Additional work will be required to replicate these results and test hypotheses consistent with this model.

Finally, we investigated whether our observed signatures of selection could be driven by adaptations of the immune system or by biological innovations in brain development. Given that polygenic selection scores do not identify individual SNPs or alleles that have experienced selection, we are limited to examine possible function of the trait-associated SNPs in bulk that contributed to observed signatures of selection. The cis-genetic component of gene expression contributes up to 30-50% of overall heritability of complex traits, but there is considerable variation among traits ^102^. For example, while cis-eQTLs contribute only 8% (s.e.=2%) to heritability of schizophrenia, they contribute up to 32% (s.e.=10%) of putamen volume ^102^. We discovered a significant enrichment of combined immune tissue eQTLs including whole blood, monocytes, and CD4+ T-cells among SNPs influencing putamen and intracranial volumes, subjective well-being, height, and type 2 diabetes (Figure 2; Supplementary Figure 5, Supplementary Table 4). We also observed an enrichment of brain tissue eQTLs among SNPs influencing schizophrenia, height, and BMI. These findings are concordant with previously reported contribution of specific cell types to heritability of complex traits (Figure 2; Supplementary Figure 5, Supplementary Table 4) ^50^. Gene set enrichment analysis indicated that the genes implicated by eQTLs in the top GWAS associations for several traits under selection cluster in biologically plausible molecular functions and biological processes, such as “neurogenesis, neuron differentiation, and neuron development” (Supplementary Table 5).

In conclusion, we demonstrate that weak polygenic adaptation, not strong selective sweeps, has influenced the evolutionary history of common genetic variation underlying neuropsychiatric disorders, behavioral traits, and brain structure volumes. The lack of evidence for strong sweeps is consistent with evolutionary scenarios in which selection coefficients for individual trait associated variants were not strong, but also consistent with hypotheses relating to fitness trade-offs between effects on multiple traits for highly polygenic phenotypes. Schizophrenia-protective alleles may have experienced both ancient and very recent positive polygenic selection, consistent with hypotheses positing shifting selection pressures between ancient and modern times due to ancestral adaptation or neutrality and fluctuating environments^103^. Furthermore, we provide genetic evidence of concerted evolution between height and total ICV, but mosaic evolution of subcortical brain structure volumes. Finally, we suggest that adaptive innovations of the immune system may also influence brain evolution among humans.

## Acknowledgements

We thank Dr. Jeremy Berg, Yair Field, and Jonathan Pritchard for helpful discussions and again thank Dr. Field for sharing scripts used to calculate standard error and p-value for the tSDS analysis. Additionally, we would like to thank Mr. Donald Hucks for assistance in manuscript preparation and Mr. Peter Straub for his assistance in data preparation. We wish to thank the individuals who participated in each study and the investigators who shared their results with the larger scientific community. Finally, we wish to acknowledge the Psychiatric Genomics Consortium, Social Science Genetic Association Consortium, Diabetes Genetics Replication and Meta-Analysis Consortium, Enhancing Neuro-Imaging Genetics through Meta-Analysis consortium, Genetic Investigation of Anthropometric Traits consortium, and the International Genomics of Alzheimer’s Project.

## Funding

This work was supported, in part, by the Conte Center for Computational Neuropsychiatric Genomics: E.R.B., E.A.K., L.K.D., and B.E.S. are supported, in part by NIH/NIMH grant 3P50MH094267-04S1. J.L.S. and P.T. are also supported, in part, by NIH grant U54 EB20403. S.E.F is supported by the Max Planck Society.

### Psychiatric Genomics Consortium TS/OCD Workgroup

We wish to acknowledge the contribution of the TS/OCD PGC Working group for sharing summary results of GWAS prior to their publication, and for their helpful reviews of this manuscript.

### International Genomics of Alzheimer’s Project

We thank the International Genomics of Alzheimer’s Project (IGAP) for providing summary results data for these analyses. The investigators within IGAP contributed to the design and implementation of IGAP and/or provided data but did not participate in analysis or writing of this report. IGAP was made possible by the generous participation of the control subjects, the patients, and their families. The i-Select chips was funded by the French National Foundation on Alzheimer’s disease and related disorders. EADI was supported by the LABEX (laboratory of excellence program investment for the future) DISTALZ grant, Inserm, Institut Pasteur de Lille, Université de Lille 2 and the Lille University Hospital. GERAD was supported by the Medical Research Council (Grant n° 503480), Alzheimer’s Research UK (Grant n° 503176), the Wellcome Trust (Grant n° 082604/2/07/Z) and German Federal Ministry of Education and Research (BMBF): Competence Network Dementia (CND) grant n° 01GI0102, 01GI0711, 01GI0420. CHARGE was partly supported by the NIH/NIA grant R01 AG033193 and the NIA AG081220 and AGES contract N01-AG-12100, the NHLBI grant R01 HL105756, the Icelandic Heart Association, and the Erasmus Medical Center and Erasmus University. ADGC was supported by the NIH/NIA grants: U01 AG032984, U24 AG021886, U01 AG016976, and the Alzheimer’s Association grant ADGC-10-196728.

## IGAP Material and methods

International Genomics of Alzheimer’s Project (IGAP) is a large two-stage study based upon genome-wide association studies (GWAS) on individuals of European ancestry. In stage 1, IGAP used genotyped and imputed data on 7,055,881 single nucleotide polymorphisms (SNPs) to meta-analyse four previously-published GWAS datasets consisting of 17,008 Alzheimer’s disease cases and 37,154 controls (The European Alzheimer’s disease Initiative – EADI the Alzheimer Disease Genetics Consortium – ADGC The Cohorts for Heart and Aging Research in Genomic Epidemiology consortium – CHARGE The Genetic and Environmental Risk in AD consortium – GERAD). In stage

2, 11,632 SNPs were genotyped and tested for association in an independent set of 8,572 Alzheimer’s disease cases and 11,312 controls. Finally, a meta-analysis was performed combining results from stages 1 & 2.

---

Supplementary Table 1. Description of GWAS studies used for 25 phenotypes. The total number of participant subjects, number of SNPs tested, lambda GC, LD score regression, and number of SNPs reaching significance at varying thresholds after PLINK clumping (R2 = 0.25, window = 500kb) are shown.

Supplementary Table 2. F3 statistic and genetic value spearman correlations. Spearman's rho for the relationship between the F3 statistic and the genetic values (Methods) of the 52 Human Genome Diversity Panel populations included in the Qx score analysis.

Supplementary Table 2. F3 statistic and genetic value spearman correlations. Spearman's rho for the relationship between the F3 statistic and the genetic values (Methods) of the 52 Human Genome Diversity Panel populations included in the Qx score analysis.

Supplementary Table 4. Tables showing eQTLs (rs id, chromosome, position, statistics associated with eQTL tests, associated eGene, and whether it is in the HLA region) among the trait-associated variants for eleven phenotypes exhibiting evidence of polygenic selection. Each phenotype and eQTL dataset (GTEx and ImmVar) are listed in a separate tab.

Supplementary Table 5. Tables listing biological processes and molecular functions enriched among the eGenes implicated by trait-associated variants in tissues and phenotypes showing significant eQTL enrichment (i.e. schizophrenia_brain, intracranial volume_immune, and subjective well-being_immune). Gene set enrichment analysis tool (http://software.broadinstitute.org/gsea/index.isp) was used to perform enrichment analysis of gene sets.

Supplementary Figure 1. Workflow for enrichment analyses. iHS and F_st_ enrichment analyses excluded Neanderthal SNPs while the NSS analysis did not. SNPs fixed in all 52 Human Genome Diversity Panel populations, and SNPs fixed in the French Reference Population were excluded for the ancient polygenic selection analysis. AF= Allele Frequencies, 1KF = 1000 Genomes Project, HLA = Human leukocyte antigen.

Supplementary Figure 2. Ancient polygenic adaptation enrichment distributions. The five brain-related phenotypes showing significant evidence of ancient polygenic adaptation are shown. The blue line represent the Qx test statistic compared to a background of 1000 permuted genetic drift models represented as histograms.

Supplementary Figure 3. F_3_ statistic correlation with genetic values from 52 Human Genome Diversity Panel populations. Populations are grouped by broad regional groups: orange = Africa, green = Oceania, pink = East Asia, purple = Americas, red = East Asia, yellow = Middle East, blue = Europe. None of these correlations are significant (see Supplementary Table 2).

Supplementary Figure 4. Recent polygenic adaptation across the genome. The five brain-related phenotypes with significant evidence of recent polygenic adaptation are shown. Each gold dot represents the mean tSDS of 1,000 SNPs. SNP bins are ranked from

GWAS p-value ~1 (left) to GWAS p-value ~0 (right). Positive tSDS corresponds to selection acting on the risk/trait-increasing allele while a negative tSDS corresponds to selection acting on the protective/trait-decreasing allele. The dashed-line represents the null expectation of no selection.

Supplementary Figure 5. eQTL enrichment in the brain and immune tissues for eleven phenotypes (including SNPs in the HLA region) exhibiting evidence of polygenic selection (significant Qx or tSDS after multiple testing correction). Light green bars represent each immune tissue or cell type: whole blood, monocytes, and cd4+ T cells, while the dark green represents enrichment in a combination of the three immune tissues. Light blue bars represent each brain tissue, while the dark blue represents enrichment in a combination of ten brain tissues, or all ten brain tissues minus cerebellum. The black dashed line represents a p-value of 0.05. The red dashed line represents the significant p-value threshold (0.004) after accounting for 13 tissues.

Supplementary Figure 6. Ancestral and derived trait-associated alleles and their frequencies. (A) Proportion of trait-associated derived alleles by minor allele frequency bins for ten neuropsychiatric disorders and three personality traits; (B) Proportion of trait associated derived alleles by minor allele frequency bins for total intracranial volume, seven subcortical brain structure volume traits, and four complex traits without neuropsychiatric associations.

